# GenoDup Pipeline: a tool to detect genome duplication using the dS-based method

**DOI:** 10.1101/384388

**Authors:** Yafei Mao, Noriyuki Satoh

## Abstract

Understanding whole genome duplication (WGD), or polyploidy, is fundamental to investigating the origin and diversification of organisms in evolutionary biology. The wealth of genomic data generated by next generation sequencing (NGS) has resulted in an urgent need for robust and accurate tools to detect WGD. Here, we present a useful and user-friendly pipeline called GenoDup for inferring WGD using the dS-based method. We have successfully applied GenoDup to identify WGD in empirical data from both plants and animals. The GenoDup Pipeline provides a reliable and useful tool to infer WGD from NGS data.

## INTRODUCTION

Whole (large-scale)-genome duplication (WGD), or polyploidy, has been regarded as an evolutionary landmark in the origins and diversification of animals, plants, and other evolutionary lineages (Van De Peer, et al. 2017). Previous studies have shown that WGD plays an important role in enhancing speciation and reducing risks of extinction. Moreover, evolutionary novelty can be generated by duplicated genes via subfunctionalization, neofunctionalization, and dosage effects under WGD (Glasauer and Neuhauss 2014; Van De Peer, et al. 2017). Therefore, identification of WGD is the first step to understanding the impacts of WGD and the fates of duplicated genes. WGD is now known to be a common event in plants, since the availability of genomic data generated by next generation sequencing (NGS). However, recent studies also suggest that WGD is a common evolutionary force in animals (Li, et al. 2018; Van De Peer, et al. 2017). Hence, an easy-to-use pipeline is urgently needed to infer WGD using NGS data.

There are three main approaches to infer WGD with NGS data (Tiley, et al. 2016). First, identification of synteny blocks is the most straightforward method to detect WGD, but it requires high-quality genome assembly, and sadly, many genomes have not yet reached that assembly quality. Second, phylogenetic analysis of gene families can unravel WGD when organisms have undergone extensive gene loss or genome shuffling, but the uncertainties of gene tree reconstruction is a serious limitation and heavy computation is required. Finally, analysis of rates of synonymous substitutions per synonymous site (dS) of duplicated genes (the dS-based method or age distribution method) is the most common and widely used approach to infer WGD.

The dS-based method is a fragmented step-wise process (Vanneste, et al. 2014). Multiple software packages are required to build gene pairs, align sequences, and calculate dS values. Usually, there are two approaches to build gene pairs using the dS-based method. First, gene pairs can be generated using gene family cluster information as paralogous gene pairs. Second, gene pairs can be generated by synteny information as anchor gene pairs. Consistent results from both approaches yield a credible conclusion.

DupPipe is a web-based method to infer WGD using the dS-based method, but users need to wait for a long time to get results (Barker, et al. 2010). In addition, DupPipe only calculates dS values based on paralogous gene pairs. Here, we develop an open-source script called GenoDup Pipeline, which integrates all processes with only one command as well as it calculates dS values based on paralogous gene pairs or/and anchor gene pairs.

## IMPLEMENTATION

### GenoDup Pipeline architecture

GenoDup Pipeline is written in Python with all alignment of sequences, building gene pairs, and dS value calculations. BioPython must be installed and three more executable dependencies are needed: MAFFT (Katoh, et al. 2002), Translatorx (Abascal, et al. 2010), and Codeml package in PAML (Yang 2007). Nuclear protein coding sequences and corresponding protein sequences are mandatory input files to run GenoDup Pipeline. In addition, gene family cluster information or anchor gene pairs information is another mandatory input file for paralogous gene pairs and anchor gene pairs approach, respectively. Once input files have been inputted appropriately, 3 subroutines in GenoDup run as follows (Figure 1).

**Figure 1.**
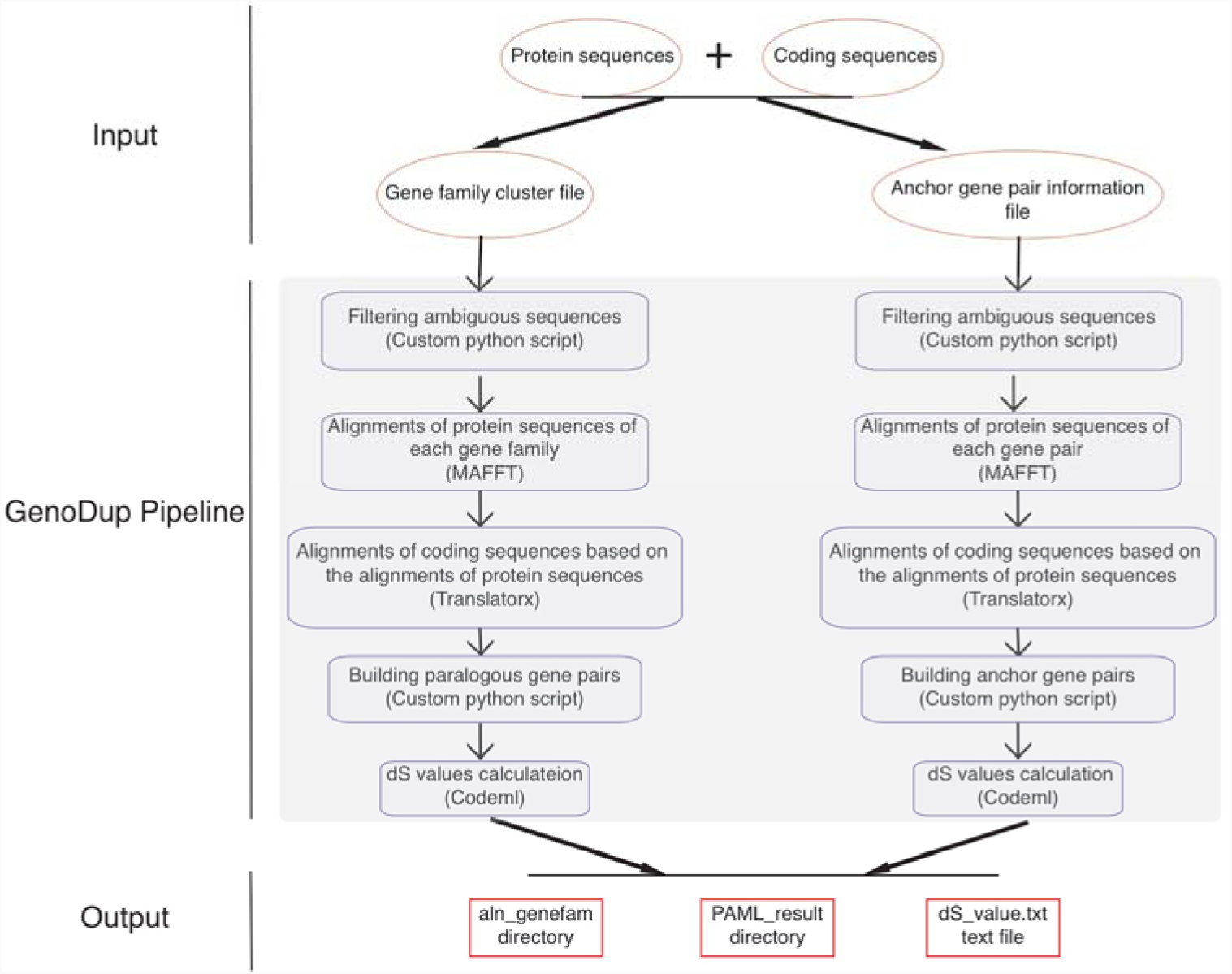
Workflow in GenoDup Pipeline. Orange oval boxes represent input files. Blue boxes represent three subroutines in the GenoDup Pipeline. Red boxes represent output files generated by GenoDup Pipeline.

(1). Alignment of gene pairs: GenoDup Pipeline can automatically align gene pairs using MAFFT and Translatorx. Before performing gene pair alignments, GenoDup first filters ambiguous sequences that contain ‘N’, and removes sequences that nuclear protein coding sequences cannot be translated to match corresponding protein sequences. Then, MAFFT is used to perform alignments of protein sequences with parameters: localpair and maxiterate: 1000; and Translatorx is used to align nuclear protein coding sequences based on alignments of the corresponding protein sequences.

(2). Building gene pairs: there are two ways to build gene pairs in GenoDup Pipeline. The first way requires a gene family cluster information file and a number (N) as input. Only the gene family, which contains less than N genes, will be used to build gene pairs. In addition, GenoDup Pipeline builds n(n-1)/2 paralogous gene pairs within a gene family (n is the number of genes in a gene family). We recommend OrthoMCL to generate gene family cluster information (Li, et al. 2003). The second way requires an anchor gene pair information file. The anchor gene pair information file contains paralogs located on same synteny blocks and we recommend MCScanX (Wang, et al. 2012) or i-ADHoRe (Proost, et al. 2011) to generate anchor gene pair information.

(3). dS value calculations: based on alignments of sequences, GenoDup can automatically build control files, required by Codeml, for each gene pair. Codeml package in PAML is used to calculate dS values with parameters: noisy = 9, verbose = 1, runmode = -2, seqtype = 1, CodonFreq = 2, model = 0, NSsites = 0, icode = 0, fix_kappa = 0, kappa = 1, fix_omega = 0, and omega = 0.5.

All of the 3 subroutines above can automatically run in the GenoDup Pipeline, which finally generates two directories and one text file as output. One directory called aln_genefam contains all nuclear protein coding sequence alignments in Fasta format. The other directory, called PAML_result contains all results generated by Codeml, and a text file called dS_value.txt contains all dS values of gene pairs. We also provided an R script (plot_GenoDup.r) to plot dS distributions.

### Empirical data validation

To evaluate the performance of the GenoDup Pipeline, we applied it to two empirical data: one is a model plant (*Arabidopsis thaliana*) and the other is a model animal (*Oncorhynchus mykiss*). *Arabidopsis thaliana* has undergone two independent WGD (alpha and beta WGD) and *Oncorhynchus mykiss* has experienced four independent WGD (Ss4R, Ts3R, and Two-rounds WGD) (Berthelot, et al. 2014; Vanneste, et al. 2014).

We downloaded the nuclear protein coding sequences, protein sequences, and genome annotation files of *Arabidopsis thaliana* from Ensembl Plants (http://plants.ensembl.org/Arabidopsis_thaliana/Info/Index). We generated gene family clusters with OrthoMCL and clustered 48,307 genes of *Arabidopsis thaliana* into 5,962 gene families. We set N as 15, meaning that the gene families containing less than 15 genes will be used to built gene pairs, and we generated 68,231 paralogous gene pairs. The entire analysis ran in 8 .8 hours with 4 cores. The result of the analysis showed an obvious peak (dS value range: 0.5∼1) in the dS distribution of paralogous gene pairs (Figure 2a). On the other hand, we generated the anchor gene pair information with MCScanX. We used MCScanX to generate 99,309 anchor gene pairs and the entire analysis ran in 76.8 hours with 4 cores (Table 1). The result of the analysis showed the same peak (dS value range: 0.5∼1) in the dS distribution of anchor gene pairs as the result based on gene family cluster information (Figure 2b).

**Table 1.** Statistics of empirical data validation in GenoDup Pipeline.

|  | *Arabidopsis thaliana* |  | *Oncorhynchus mykiss* |  |
| --- | --- | --- | --- | --- |
|  | Paralogous gene pairs | Anchor gene pairs | Paralogous gene pairs | Anchor gene pairs |
| The number of gene pairs | 68,231 | 99,309 | 42,888 | 26,880 |
| Running Time (h) | 8.88 | 76.8 | 7.63 | 17.65 |

**Figure 2.**
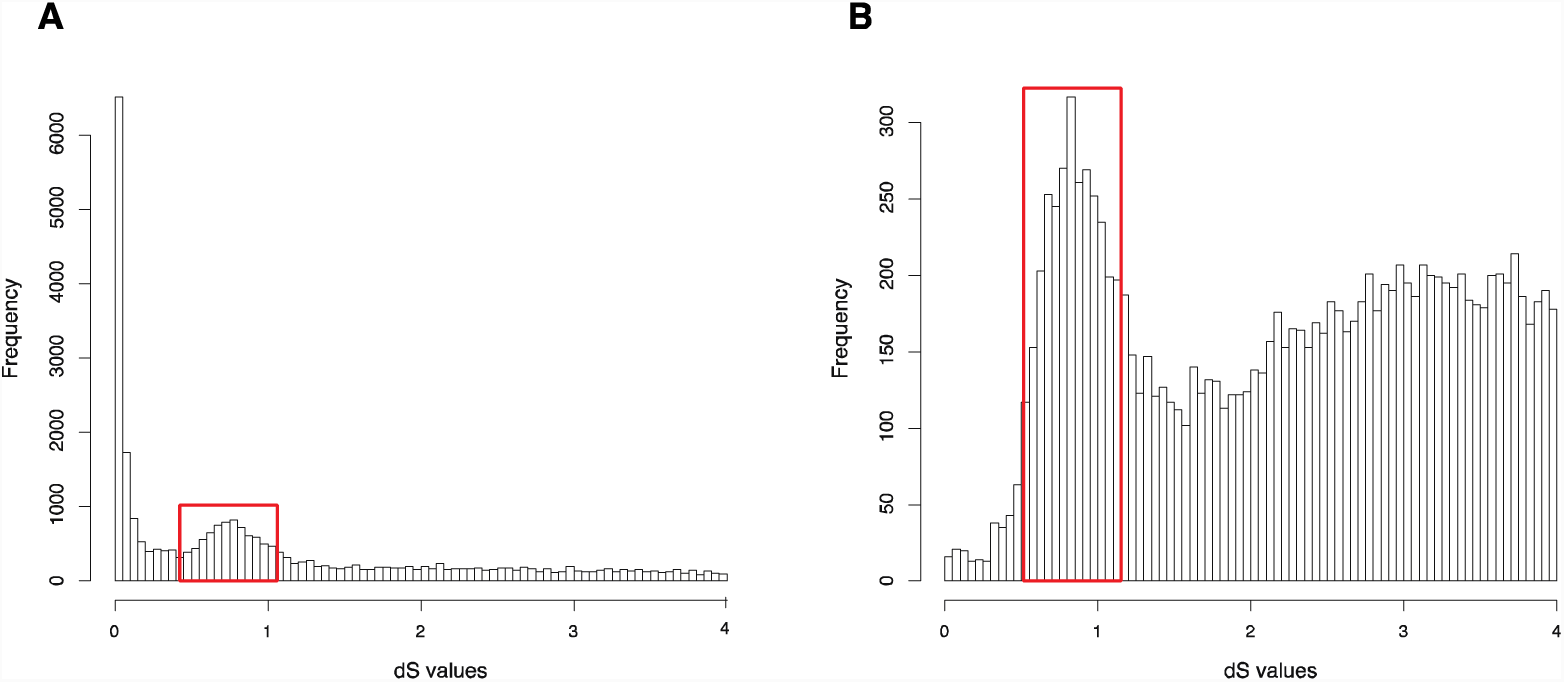
dS distributions of paralogous gene pairs and anchor gene pairs in *Arabidopsis thaliana*. (A). The peak (dS value range: 0.5∼1) marked with a red box represents a signal as alpha WGD of *Arabidopsis thaliana* in the dS distributions of paralogous gene pairs. (B). The peak (dS value range: 0.5∼1) marked with a red box represents a signal as alpha WGD of *Arabidopsis thaliana* in the dS distributions of anchor gene pairs.

We downloaded the nuclear protein coding sequences, protein sequences, and genome annotation files of *Oncorhynchus mykiss* from GENEOSCOPE (http://www.genoscope.cns.fr/trout/data/). We generated gene family cluster by OrthoMCL and clustered 46,585 genes of *Oncorhynchus mykiss* into 6,562 gene families. We set N as 15, meaning that the gene families containing less than 15 genes will be used to built gene pairs, and we generated 42,888 paralogous gene pairs. The entire analysis ran in 7.63 hours with 4 cores. The result of the analysis showed two obvious peaks (dS value ranges: 0.1∼0.5 and 1.2∼2) in the dS distribution of paralogous gene pairs (Figure 3a). On the other hand, we generated anchor gene pair information identified by MCScanX. We used MCScanX to generate 26,880 anchor gene pairs and the entire analysis ran in 17.65 hours with 4 cores (Table 1). The result of the analysis showed the same peak (dS value range: 0.1∼0.5) in the dS distribution of anchor gene pairs as the result based on gene family cluster information (Figure 3b).

**Figure 3.**
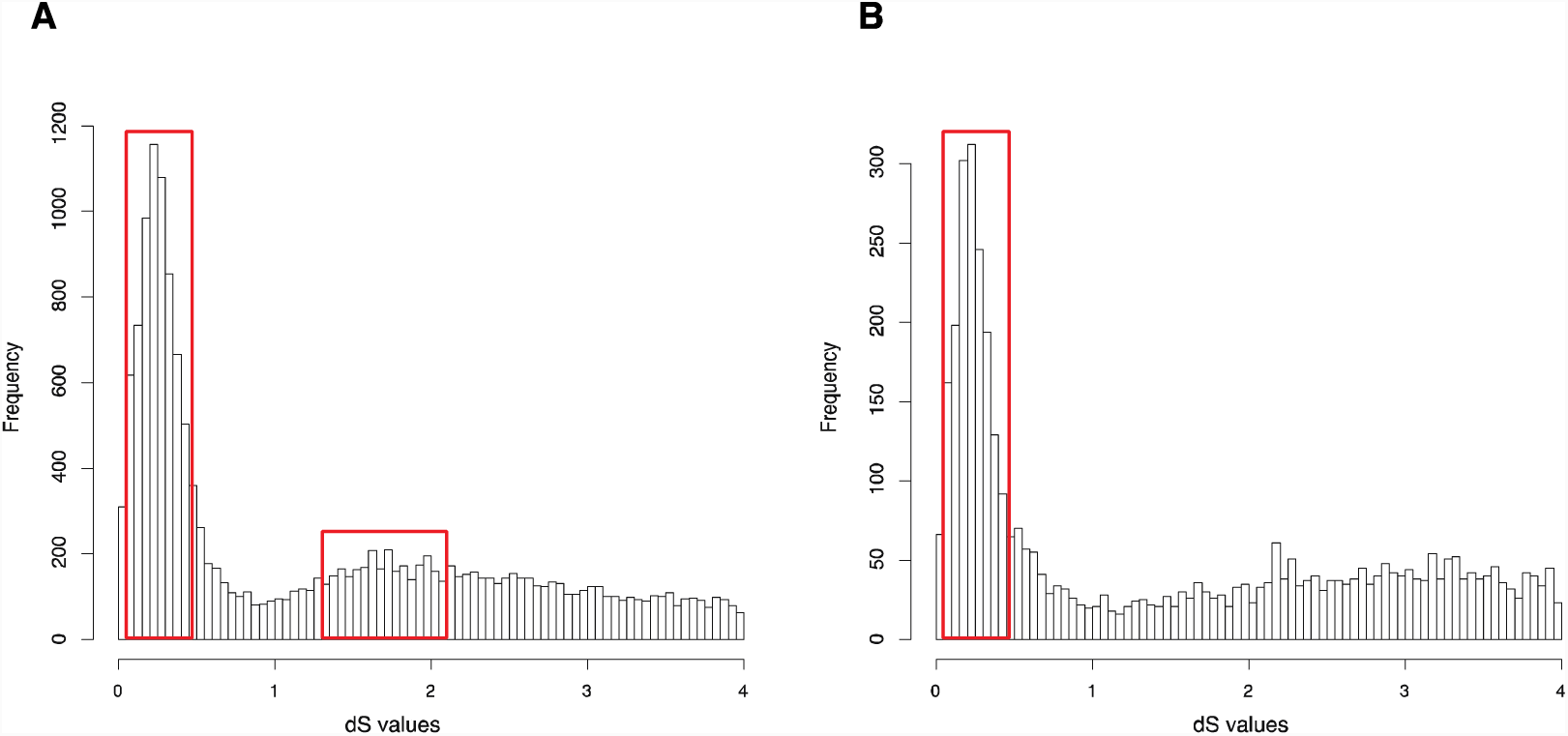
dS distributions of paralogous gene pairs and anchor gene pairs in *Oncorhynchus mykiss*. (A). The peaks (dS value ranges: 0.1∼0.5 and 1.2∼2) marked with red boxes represent signals of the Ss4R and Ts3R WGDs of *Oncorhynchus mykiss*, respectively, in the dS distributions of paralogous gene pairs. (B). The peak (dS value range: 0.1∼0.5) marked with a red box represents the signal attributable to Ss4R WGD of *Oncorhynchus mykiss* in the dS distributions of anchor gene pairs.

Our GenoDup Pipeline represented consistent result as the previous study on *Arabidopsis thaliana* and *Oncorhynchus mykiss*, respectively (Berthelot, et al. 2014; Vanneste, et al. 2014), indicating that GenoDup Pipeline is a reliable tool to infer WGD using the dS-based method.

## DISCUSSION

The rapid development of NGS technologies has enabled generation of massive amounts of data, allowing us to better understand the evolutionary history of all organisms. WGD is suggested to have occurred in diverse organismal groups (Li, et al. 2018; Van De Peer, et al. 2017); thus, a reliable, efficient tool to detect WGD with NGS data is greatly needed. We developed a reliable and easy-to-use tool called GenoDup Pipeline to infer WGD using the dS-based method. GenoDup Pipeline is written by Python and can be run with one command. It is easy to use for researchers who have a little background in bioinformatics.

In GenoDup Pipeline, it is faster to run to take gene family cluster information as input rather than to take anchor gene pair information as input. Because GenoDup Pipeline aligns sequences for each gene family when taking gene family cluster information whereas GenoDup Pipeline aligns sequences for anchor gene pairs individually when taking anchor gene pair information as input. Alignments of anchor gene pairs individually cost much time in GenoDup Pipeline.

Our empirical validation shows that the analysis of *Arabidopsis thaliana* generated with GenoDup Pipeline presented an obvious signal for one WGD event (alpha WGD) but the signal of the second WGD event (beta WGD) was lost. This result is consistent with previous studies because the dS methods cannot infer WGD when organisms have undergone extensive gene loss or genome shuffling, especially for plants (Rabier, et al. 2014; Tiley, et al. 2016). Additionally, the *Oncorhynchus mykiss* analysis with GenoDup showed two obvious two WGD signals (Ss4R and Ts3R) in the dS distribution of paralogous gene pairs because of high conservation in the *Oncorhynchus mykiss* genome (Rabier, et al. 2014). Yet, The Ts3R signal was lost in the distribution of anchor gene pairs because there were few anchor gene pairs in the analysis (Table 1). Moreover, the both analysis on *Oncorhynchus mykiss* do not present the Two-round WGDs because it is too ancient to infer with the dS-based method. Importantly, the dS-based method has been debated to generate artificial signals, because of dS saturation when the dS value >1 (Vanneste, et al. 2012). Thus, GenoDup Pipeline is suitable to infer WGD when the dS value <1, and other evidence from phylogenetic analysis and/or synteny blocks analysis is needed to draw a conclusive result.

In all, with more and increasingly large sequencing projects (1KP, 10KP-EBP, 1KITE, and i5K) developing (Li, et al. 2018; Van De Peer, et al. 2017), we present a reliable and user-friendly tool to infer WGD using the dS-based method.

## ACKNOWLEDGEMENTS

This study was supported by OIST and was funded by JSPS grant (No. 17J00557 to YFM). We thank Dr. Steven D. Aird for editing the manuscript.

## Availability and requirements

Project name: GenoDup Pipeline

Source code: https://github.com/MaoYafei/GenoDup-Pipeline

Documentation: https://github.com/MaoYafei/GenoDup-Pipeline/blob/master/README.md

Operating system: Linux (Red Hat, CentOS, or Ubuntu) Programming language: Bash and Python

Other requirements: MAFFT, Translatorx, Codeml package in PAML

## Author Contributions

Yafei Mao conceived and designed the experiments, performed the experiments, analyzed the data, wrote the paper, and reviewed drafts of the paper.

Noriyuki Satoh supervised the project, wrote the paper, and reviewed drafts of the paper.

## Data Availability

The following information was supplied regarding data availability: Github: https://github.com/MaoYafei/GenoDup-Pipeline

